## Supplementary material for "Reciprocal Priming between Receptor Tyrosine Kinases at Recycling Endosomes Orchestrates Cellular Signalling Outputs": SI

### Appendix

#### TABLE OF CONTENT

Appendix Figures and Legends S1-S7

### APPENDIX FIGURES AND LEGENDS

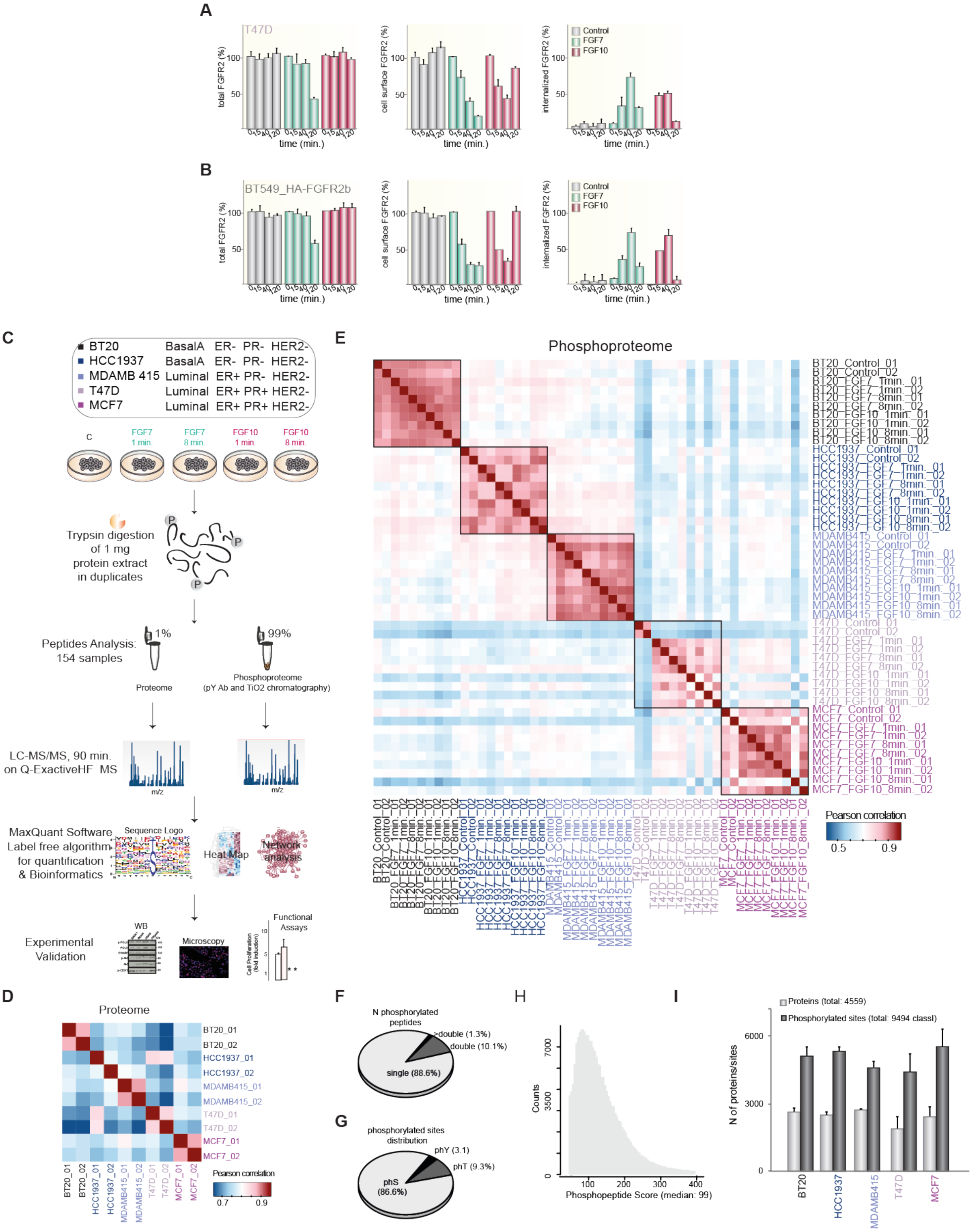


**Appendix Figure S1. Quality Assessment of TPA1 Showed Good Reproducibility.** The presence (total), internalization (internalized), and recycling (cell surface) of endogenous FGFR2 in T47D (A) or HA-FGFR2b transfected in BT549 (B) upon stimulation with FGF7 (dark green) or FGF10 (burgundy) for different time periods were quantified as described (Francavilla *et al*, 2016). Values represent the median ± SD of N=3. T47D representative pictures are shown in Figure 1B. (C) Experimental design of

TPA1. We analyzed four independent experiments (two each at either 1 or 8 min. stimulation with each FGF. This represents early signaling. (D) Heatmap of the Pearson's correlation of the whole proteome of the analyzed breast cancer cell lines showed very good reproducibility among cell lines with a Pearson correlation coefficient between 0.7 and 0.9. (E) Heatmap of the Pearson's correlation of the phosphoproteome of the stimulated breast cancer cell lines showed very good reproducibility within each cell line (Pearson correlation coefficient higher than or equal to 0.75) and differences among cell lines (Pearson correlation coefficient smaller than 0.75). (F) Distribution of phosphorylated peptides with one, two or more phosphorylated sites. (G) Distribution of identified serine (pS), threonine (pT), and tyrosine (pY) phosphorylated sites. (H) The distribution of phosphorylated peptides score showed that most of the peptides were identified with high Andromeda score (median: 99). (I) Number of identified proteins and phosphorylated sites in each breast cancer cell line.

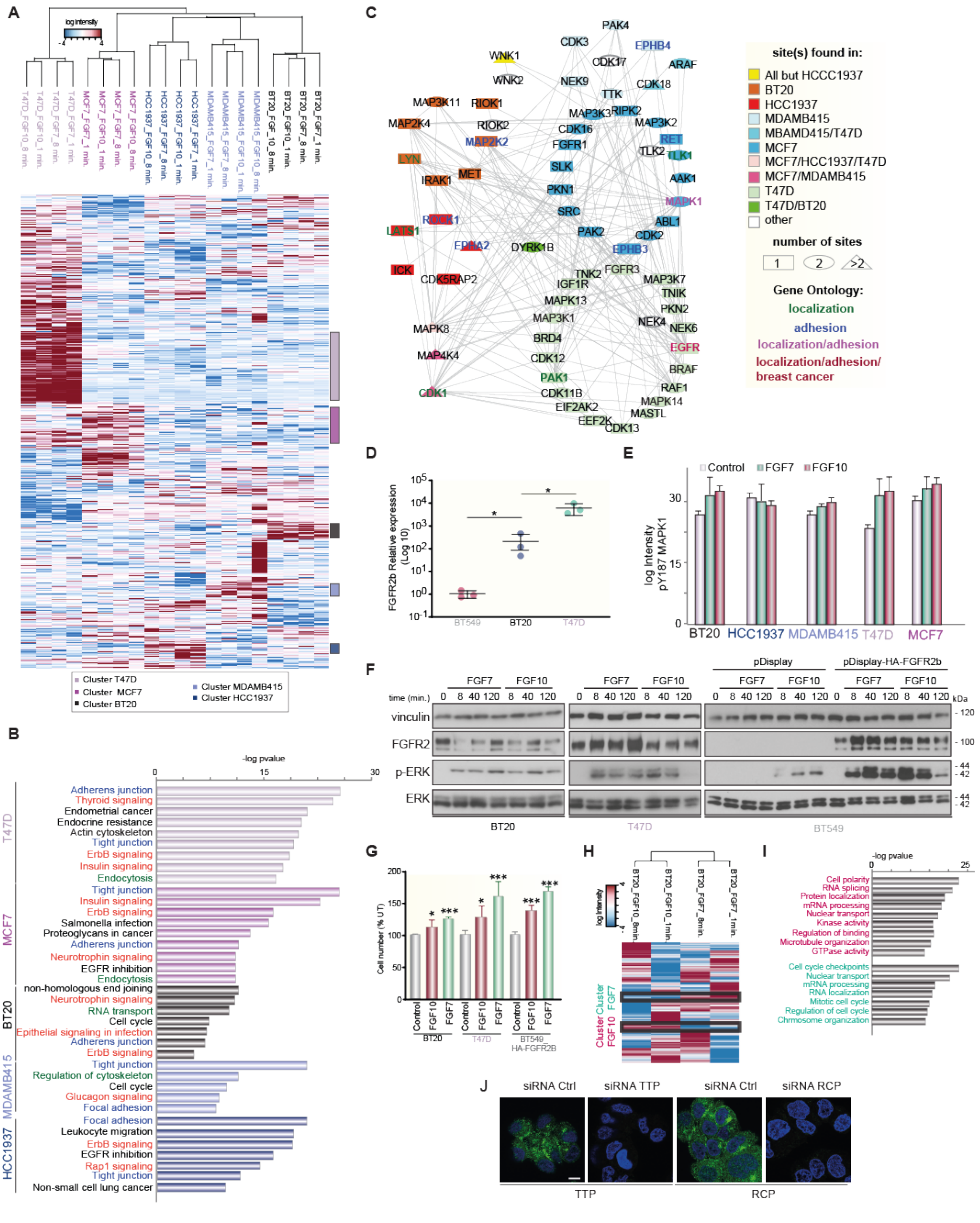


**Appendix Figure S2. TPA1 Uncovers Common and Distinct Signaling Pathways upon FGFR2b Stimulation in breast cancer Cells.** (A) The hierarchical clustering of the phosphorylated sites identified in five breast cancer cell lines upon stimulation with FGF7 and FGF10 for short time points (1 and 8 min.) showed distinct clusters representing proximal FGFR2b signaling for each cell line. Each cluster is represented by a colored bar on the right. The intensity of phosphorylated sites is presented on the logarithmic scale with intensity below and above the mean color-coded in blue and red, respectively. (B) KEGG (Kyoto Encyclopedia of Genes and Genomes) pathways enriched in each of the five clusters shown in A. KEGG terms related to “adhesion” are color-coded in dark blue; terms related to “establishment of localization” are color-coded in dark green; terms related to “signaling pathways” are color-coded in orange. All these terms with the exception of “establishment of localization” in the cell line HCC1937 were enriched in all the analyzed cell lines upon FGFR2b activation. (C) Network of phosphorylated kinases belonging to the Gene Ontology (GO) categories “establishment of localization”, “adhesion”, or found associated to breast cancer according to the database DISEASES (Pletscher-Frankild *et al*, 2015). The network was based on STRING and visualized with Cytoscape. The nodes are color-coded based on the cluster the first identified phosphorylated site was found in. Different shapes represent the number of sites found on the phosphorylated protein. The color of the text is based on the GO categories as in B and on the DISEASES database. (D) Relative mRNA expression of Fgfr2 splicing variant “b” in the indicated breast cancer cell lines as determined by qPCR. (E) Quantification of MAPK1 Y187-containing phosphorylated peptide identified in the dataset shown in (A) showed differences upon FGF7 or FGF10 stimulation over untreated cells. Values are in log scale and are the median ± SD of two (control conditions) or four biological replicates (stimulated conditions). (F) Lysates from BT20, T47D or BT549 (either transfected with pDisplay_HA-FGFR2b or pDisplay empty vector) cell lines stimulated with FGF7 or FGF10 for the indicated time points or left unstimulated (0 min.) were immunoblotted with the indicated antibodies. The phosphorylation of ERK was weak and sustained in BT20 upon both FGF7 and FGF10 stimulation. We observed sustained and transient ERK activation upon FGF7 and FGF10 stimulation, respectively in both T47D and BT549 which overexpressed HA-FGFR2b. These data suggest that T47D and HA-FGFR2b-transfected BT549 respond very similarly to both FGFR2b ligands, whereas BT20 and T47D may differ in their prolonged response to FGF10 treatment. However, all three cell lines respond in a comparable manner to FGF10 stimulation up to 40 min.. Finally, the data shown in E and F show good reproducibility of MAPK/ERK phosphorylation, therefore validating our MS-based approach to study cellular signaling (Francavilla *et al*, 2013). (G) Cell number at 72 hours of BT20, T47D, and BT549 transfected with HA-FGFR2b stimulated with FGF7 (dark green) or FGF10 (burgundy). Data represent the mean ± SD of 3 experiments compared to control cells (UT). pvalue = <0.05 *, <0.01 **, <0.001*** (Student’s t-test). (H) Hierarchical clustering of the phosphorylated sites differentially quantified in BT20 stimulated with FGF7 (green) or FGF10 (burgundy). Specific clusters are highlighted with black lines. The intensity of phosphorylated sites is presented on the logarithmic scale with intensity below and above the mean color-coded in blue and red, respectively. (I) Enriched terms in the selected clusters in H. (J) T47D cells were depleted or not of TTP or RCP as indicated and the expression of TTP and RCP was checked by immunofluorescence. Bar, 5 μm.

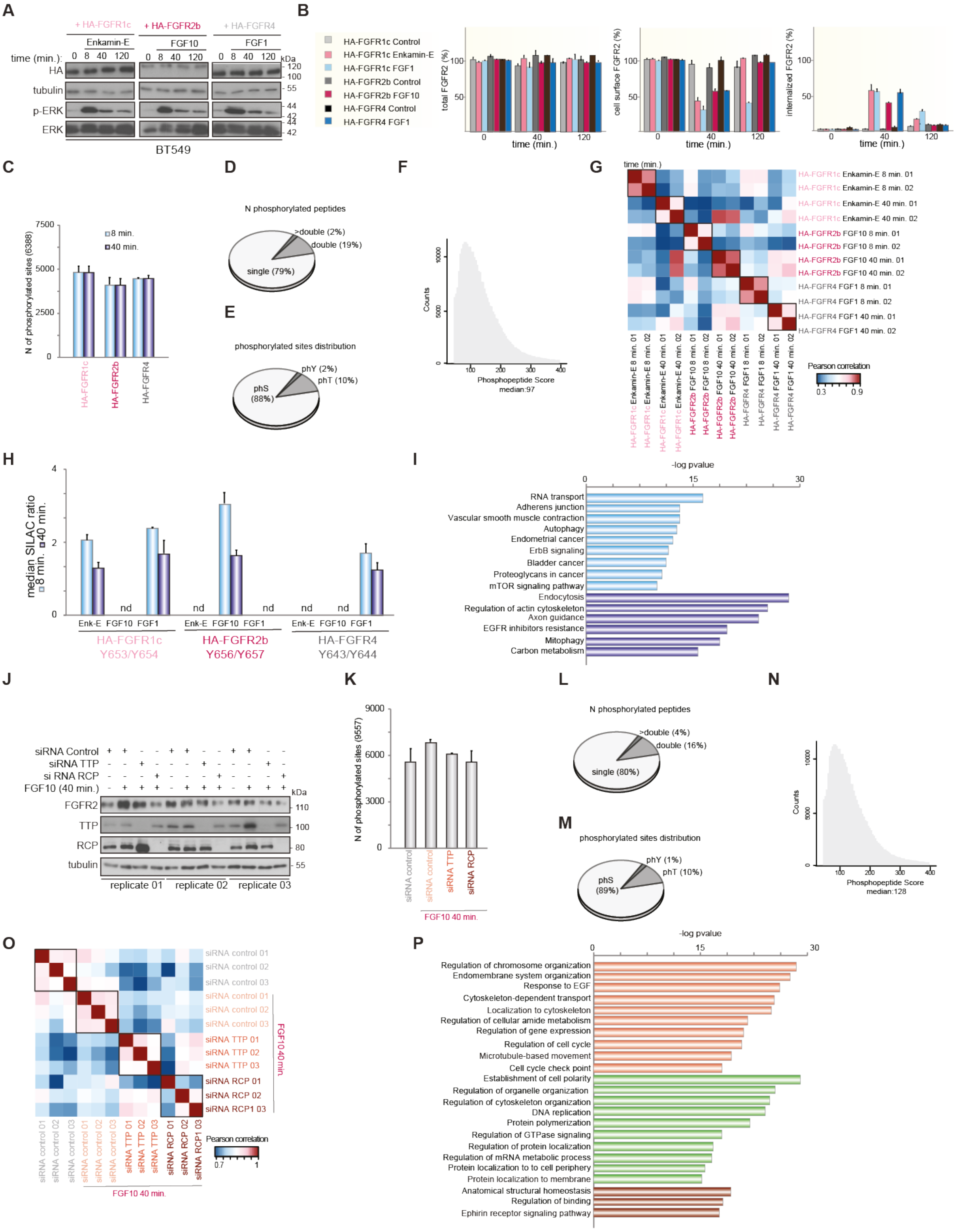


**Appendix Figure S3. Quality Assessment of TPA2 and TPA3 Showed Good Reproducibility.** (A) Lysates from BT549 transfected with HA-FGFR1c, HA-FGFR2b, or HA-FGFR4 and stimulated for different time periods with Enkamin-E, FGF10, or FGF1, respectively were immunoblotted with the indicated antibodies. We observed phosphorylation of ERK in all conditions, indicating signaling activation. (B) The presence (total), internalization (internalized), and recycling (cell surface) of transfected HA-FGFRs in BT549 upon stimulation with the indicated ligands for different time periods were quantified as described (Francavilla *et al.*, 2016). Values represent the median ± SD of three independent experiments expressed in percentage. HA-FGFR1c recycled to plasma membrane and was degraded upon Enkamin-E and FGF1, respectively, as previously shown (Francavilla *et al*, 2009). HA-FGFR2b recycled to plasma membrane upon FGF10 stimulation, confirming the data shown in Figure 1. HA-FGFR4 recycled to plasma membrane upon FGF1 stimulation, as previously shown (Haugsten *et al*, 2005). (C) Number of identified phosphorylated sites in BT549 transfected with HA-FGFR1c, HA-FGFR2b, or HA-FGFR4 and stimulated for 8 or 40 min. with Enkamin-E, FGF10, or FGF1, respectively. (D) Distribution of phosphorylated peptides with one, two or more phosphorylated sites. (E) Distribution of identified serine (pS), threonine (pT), and tyrosine (pY) phosphorylated sites. (F) The distribution of phosphorylated peptides score showed that most of the peptides were identified with high Andromeda score (median: 97). (G) Heatmap of the Pearson's correlation of the phosphoproteome of BT549 transfected with HA-FGFR1c, HA-FGFR2b, or HA-FGFR4 and stimulated for 8 or 40 min. with Enkamin-E, FGF10, or FGF1, respectively showed very good reproducibility among biological replicates (Pearson correlation coefficient higher than or equal to 0.75) and differences among stimulated cells expressing different HA-FGFRs (Pearson correlation coefficient smaller than 0.6). (H) Quantitation of the catalytic tyrosine-containing phosphorylated peptides of each transfected receptor (Y653/Y654-containing peptide on HA-FGFR1c; Y656/Y657-containing peptide on HA-FGFR2b; and Y643/Y644-containing peptide on HA-FGFR4) validated our approach based on MS-driven quantitative phosphoproteomics (Dataset EV3) to study receptor recycling-dependent signaling in transfected BT549 cells. nd: not detected. We identified the catalytic tyrosine-containing phosphorylated peptides of each transfected receptor specifically in cells transfected with that receptor, indicating the lack of auto phosphorylation in transiently transfected BT549 cells and the specificity of our MS analysis. (I) KEGG terms enriched in the clusters. Early Signaling and Recycling Receptors shown in Figure 2B are color-coded in light and medium blue, respectively. (J) Lysates from T47D depleted or not of TTP, RCP and stimulated with FGF10 for 40 min. were immunoblotted with the indicated antibodies. Replicates 01-03 refer to the biological replicates of the experiment described in Figure 2D-F. We observed reproducible depletion of TTP and RCP upon siRNA treatment. (L) Number of identified phosphorylated sites in T47D depleted of TTP, RCP or left untreated and stimulated with FGF10 for 40 min.. (K) Distribution of phosphorylated peptides with one, two or more phosphorylated sites. (M) Distribution of identified serine (pS), threonine (pT), and tyrosine (pY) phosphorylated sites. (N) The distribution of phosphorylated peptides score showed that most of the peptides were identified with high Andromeda score (median: 128). (O) Heatmap of the Pearson's correlation of the phosphoproteome of T47D depleted of TTP, RCP or left untreated and stimulated with FGF10 for 40 min. showed very good reproducibility among biological replicates (Pearson correlation coefficient higher than or equal to 0.85) and differences among stimulated T47D depleted of either TTP or RCP (Pearson correlation coefficient smaller than 0.7). (P) BP GO terms enriched in the three clusters TTP Adaptor, RCP Adaptor or Recycling Adaptors shown in Figure 2E are color-coded in orange, brown, and light green, respectively.

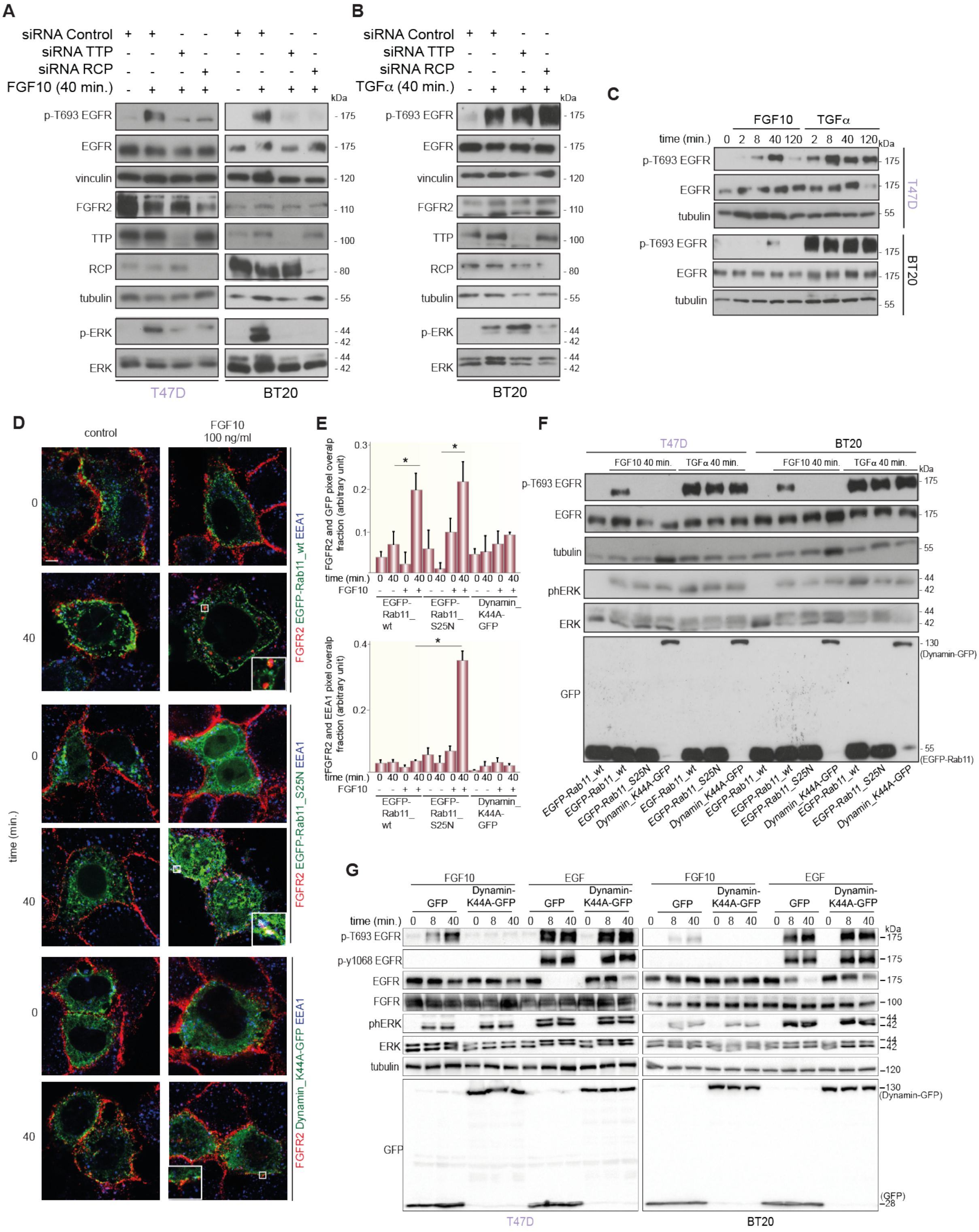

**Appendix Figure S4. FGF10- but not TGFα-induced EGFR_T693 Phosphorylation Depends on FGFR2b Recycling.** Lysates from T47D (A) and BT20 (B) left untreated or depleted of TTP or of RCP using a pool of siRNAs and stimulated or not with FGF10 or TGFα for 40 min. and from T47D and BT20 (C) stimulated or not with FGF10 or TGFα for different time periods were immunoblotted with the indicated antibodies. (D) Co-localization of FGFR2 (red) with EGFP-Rab11, EGFP-Rab11_S25N (dominant negative Rab11), or Dynamin_K44A-GFP (dominant negative Dynamin) (green) and the early endosomes marker EEA1 (blue) in T47D stimulated with FGF10 for 40 min.. Scale bar, 5 μm. (E) Red and green pixels overlap fraction (above) representing the co-localization of FGFR2 with GFP-tagged proteins (green pixels) and red and far red pixels overlap fraction (below) representing the co-localization of FGFR2 (red pixel) with the early endosomes marker EEA1 (far red pixels) in T47D stimulated with FGF10 for 40 min. Values represent the median ± SD of at least three independent experiments. Representative pictures are shown in D. *, p value<0.005 (Student's t-test). FGF10 induced the co-localization of FGFR2b (red pixels) with Rab11 (green pixels) in both wild type- and dominant negative-Rab11 transfected cells, but not the co-localization with dominant negative dynamin (green pixels) (graph above). FGF10 induced the co-localization of FGFR2b (red pixels) with EEA1 (far red pixels) in EGFP-Rab11_S25N- but not in EGFP-Rab11- or Dynamin_K44A-GFP-expressing cells (graph below). All together, these findings indicate that FGFR2b is in recycling endosomes in T47D cells transfected with EGFP-Rab11, confirming the results shown in Figure 1H-I, whereas FGFR2b is in the early endosome compartment in the presence of dominant negative Rab11, as previously shown for FGFR1 (Francavilla *et al.*, 2009). Finally, FGF10-stimulated FGFR2b is at plasma membrane in cells transfected with dominant negative Dynamin (last panel on the right in comparison with the second and the fourth panels on the right). (F, G) Lysates from T47D and BT20 transfected with EGFP-Rab11, EGFP-Rab11_S25N (dominant negative Rab11), or Dynamin_K44A-GFP (dominant negative Dynamin) and stimulated or not with FGF10 or TGFα or EGF for either 8 or 40 min. as indicated, were immunoblotted with the indicated antibodies.

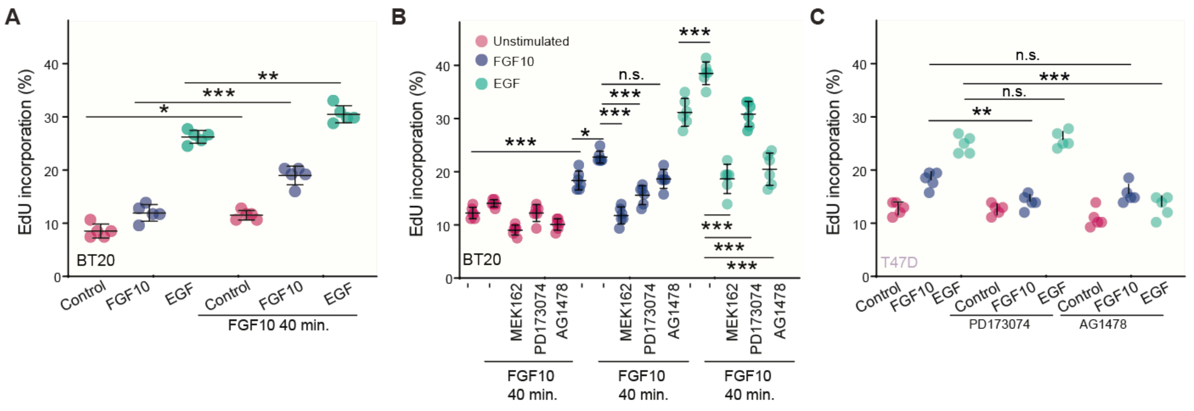


**Appendix Figure S5. Pretreatment with FGF10 Increases EGF-induced Cell Cycle Progression via FGFR, EGFR, and ERK Activation.** Percentage of EdU incorporation in BT20 pre-treated or not with FGF10 for 40 min. and stimulated or not with FGF10 or EGF (A) or incubated with MEK162, PD173074, or AG1478 before pretreatment (B). (C) Percentage of EdU incorporation in T47D stimulated with FGF10 or EGF in the presence or absence of PD173074 or AG1478 without pretreatment of FGF10 for 40 min. Data represent n=5 experiments. Significance was calculated by one-way ANOVA with Tukey test p=<0.05 *, <0.01 **, <0.001 ***.

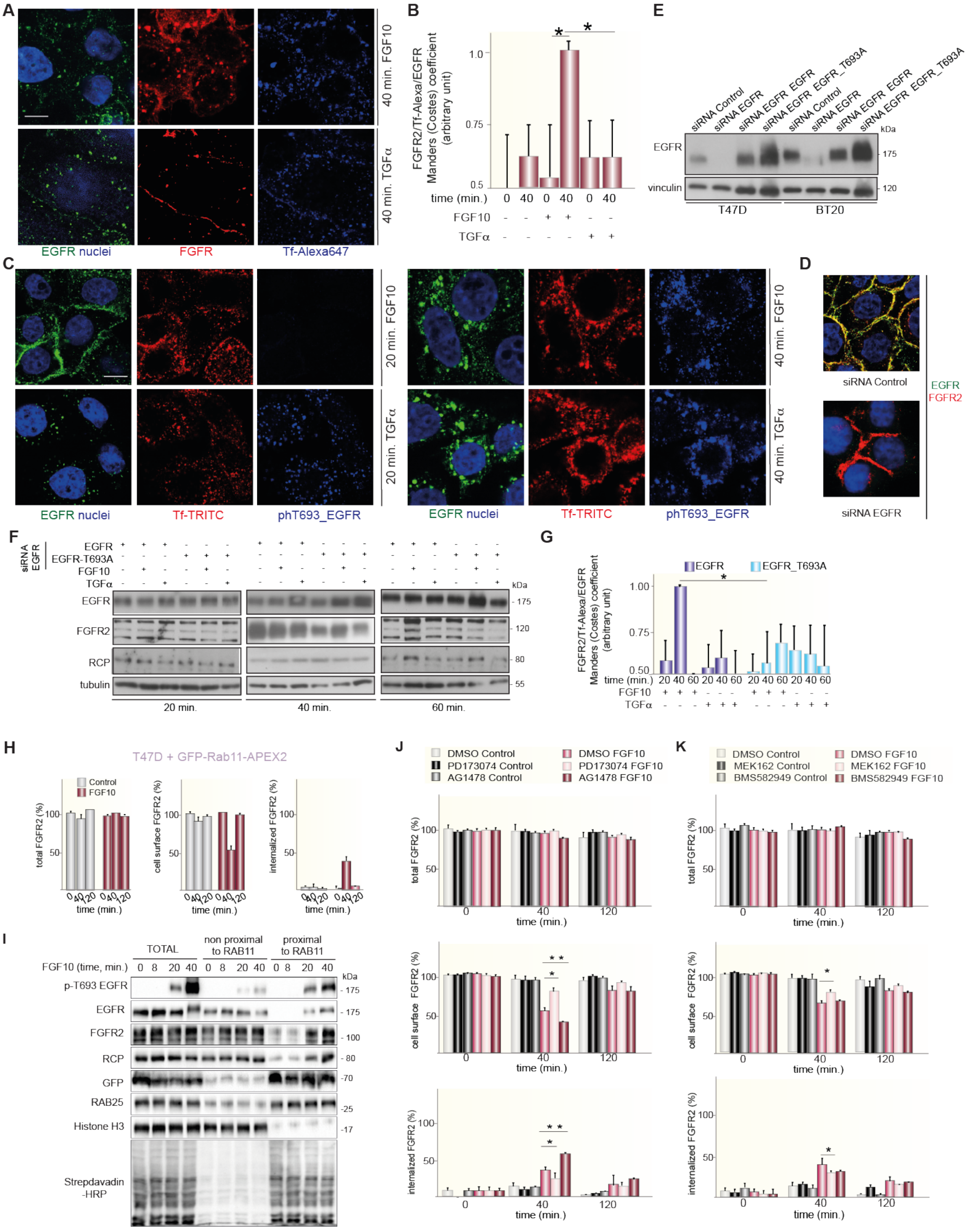

**Appendix Figure S6. FGFR2b Recycling Depends on EGFR_T693 Phosphorylation, FGFR and ERK Activation.** (A) Co-localization of FGFR2 (red), EGFR (green), and the recycling marker Tf (blue) in T47D stimulated with FGF10 or TGFα for 40 min. Scale bar, 5 μm. Related to Figure 5a. The separated channels clearly indicate the different FGFR2b localization in FGF10 or TGFα-stimulated cells. (B) Our Manders Colocalization Coefficient adapted for three channels Red, Far red and Green representing the co-localization of the combined FGFR2b and EGFR, and the recycling marker Tf in T47D stimulated for 40 min. Values represent the median ± SD of at least three independent experiments. Representative pictures are shown in Figure 5A and Figure S6A. *, p value<0.005

(Student's t-test). The Manders (Costes thresholds) coefficient confirms the co-localization of FGFR2b, EGFR, and Tf upon stimulation with FGF10 but not TGFα for 40 min. as shown in Figure 5A. (C) Co-localization of EGFR (green), phosphorylated EGFR on T693 (blue), and the recycling marker Tf (red) in T47D stimulated or not with FGF10 or TGFα for either 20 or 40 min., as indicated. Scale bar, 5 μm. Related to Figure 5C. The separated channels clearly indicate that EGFR is phosphorylated on T693 upon both FGF10 and TGFα stimulation in T47D cells. Furthermore, these images confirmed that EGFR was detected in recycling endosomes upon 40 min. stimulation with FGF10. (D) Immunofluorescence in T47D depleted or not of EGFR by siRNA showed the co-localization of EGFR (green) and FGFR2 (red) in untreated cells (left) and the lack of EGFR (green) in depleted cells (right). Scale bar, 5 μm. Lysates from T47D and BT20 cells left untreated or depleted of EGFR by siRNA followed by transfection with wt or T693A (E) and from T47D depleted of EGFR by siRNA followed by transfection with wt or T693A and stimulated or not with either FGF10 or TGFα for the indicated time periods (F) were immunoblotted with the indicated antibodies. Input samples related to Figure 5G (E) The depletion of EGFR followed by transfection with wt or T693A was efficient in both T47D and BT20 (G) Red, far red and green Manders colocalization coefficient representing the co-localization of FGFR2b, EGFR, and the recycling marker Tf in T47D depleted of EGFR by siRNA followed by transfection with wt or T693A and stimulated or not with either FGF10 or TGFα for the indicated time periods. Values represent the median ± SD of at least three independent experiments. Representative pictures are shown in Figure 5F. *, p value<0.005 (Student's t-test). Our three channel Manders colocalization coefficient (Costes thresholds) confirms the co-localization of FGFR2b, EGFR, and Tf upon stimulation with FGF10, but not TGFα, for 40 min. in wt- but not T693A-expressing cells. In T693A-expressing cells both FGFR2b and EGFR were detected at the plasma membrane in cells stimulated with both FGF10 and TGFα for all the considered time points. Upon FGF10 stimulation FGFR2b was detected in recycling endosomes at 20 but not 60 min. in wt- expressing cells, whereas EGFR was detected at plasma membrane at both 20 and 60 min.. In TGFα stimulated cells expressing wt EGFR was detected in recycling endosomes at 40 min. and at plasma membrane at 20 and 60 min., whereas FGFR2b was detected at plasma membrane at all considered time points. (I) T47D transfected with GFP-Rab11-APEX2 were stimulated with FGF10 at indicated timepoints. Biotinylation was performed followed by streptavidin bead pulldown, running the supernatant against the lysates extracted off the beads and immunoblotted with the indicated antibodies, including knownRab11 interactors RAB25 and RCP and nuclear control Histone H3. (H, J, K) The presence (total), internalization (internalized), and recycling (cell surface) of FGFR2b in T47D either transfected with GFP-Rab11-APEX (G) or treated with DMSO, PD173074, AG1478, MEK162, and BMS582949 (J, K) and stimulated with FGF10 for different time periods were quantified as described (Francavilla *et al.*, 2016). Values represent the median ± SD of three independent experiments expressed in percentage. The transfection with GFP-Rab11-APEX did not alter FGFR2b trafficking (see also Figure 1B). PD173074 and MEK162 treatment resulted in increased FGFR2b at plasma membrane and decreased FGFR2b in the cytoplasm at 40 min. stimulation, suggesting faster recycling of FGFR2b, whereas AG1478 treatment showed the opposite effect. Finally, p38 inhibition did not affect the kinetic of FGFR2b recycling.

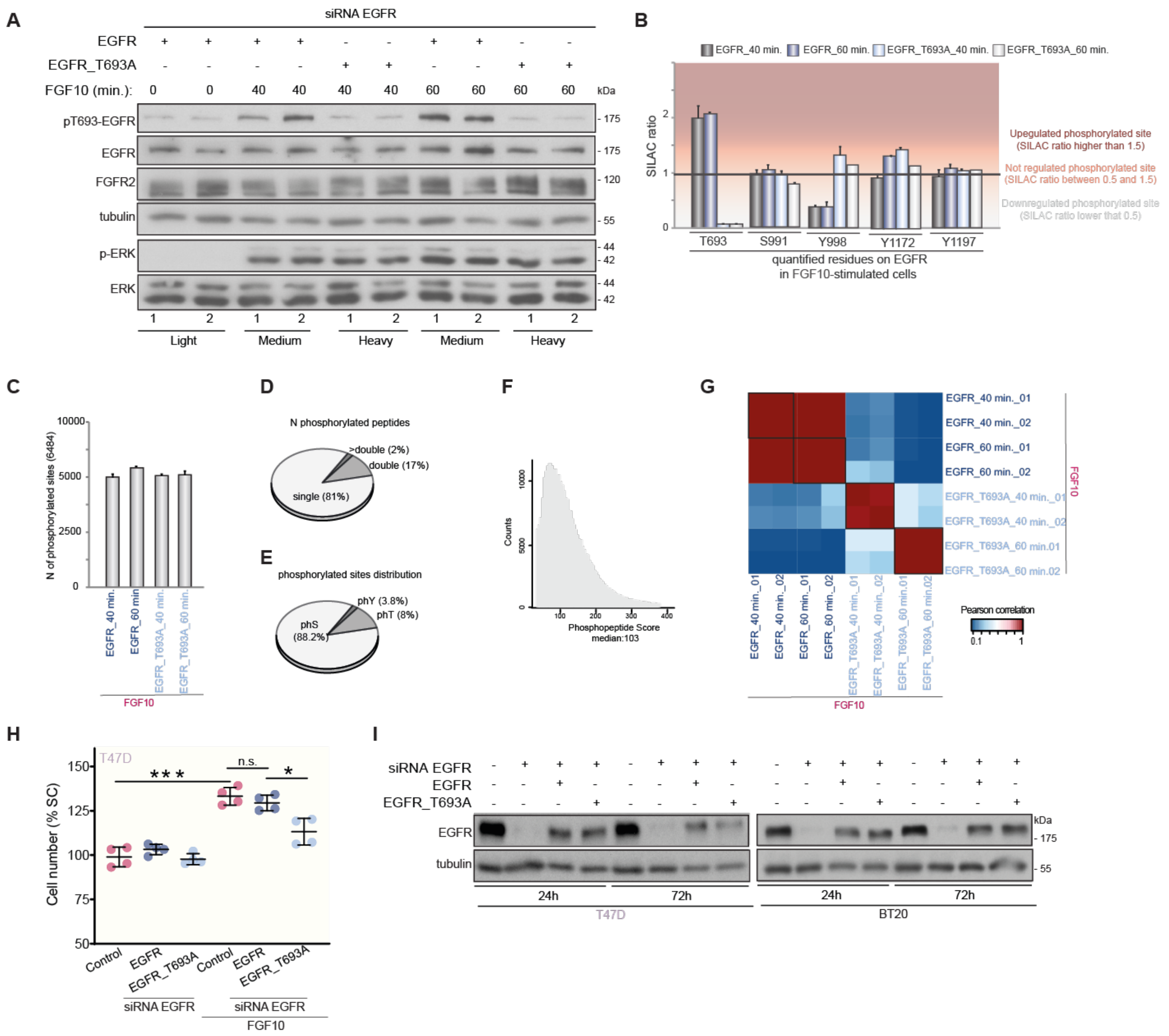


**Appendix Figure S7. Quality Assessment of the EGFR_T693 Phosphoproteomics Dataset Showed Good Reproducibility.** (A) Lysates from T47D depleted of EGFR, transfected with wt or T693A, and stimulated or not with FGF10 for 40 or 60 min. were immunoblotted with the indicated antibodies. Light, Medium, and Heavy refer to the SILAC condition and the numbers 1 and 2 to the two biological replicates. We observed reproducible transfection of EGFR constructs, phosphorylation of T693 only in wt-expressing cells, and phosphorylation of ERK in all tested conditions. (B) Quantification of EGFR phosphorylated sites identified in the dataset shown in Figure 6A showed that FGF10 induced the phosphorylation of T693 (SILAC ratio higher than 1.5) but not of other sites (SILAC ratio lower than 1.5) on EGFR. T693 was not phosphorylated in EGFR_T693A-expressing T47D. The phosphorylation of the other identified residues was not affected by the T693A mutation, with the exception of tyrosine 998 phosphorylation which increased. Values represent the median ± SD of two biological replicates. (C) Number of identified phosphorylated sites in each experimental condition. (D) Distribution of phosphorylated peptides with one, two or more phosphorylated sites. (E) Distribution of identified serine (pS), threonine (pT), and tyrosine (pY) phosphorylated sites. (F) The distribution of phosphorylated peptides score showed that most of the peptides were identified with high Andromeda score (median: 103). (G) Heatmap of the Pearson's correlation of the phosphoproteome of T47D depleted of EGFR, transfected with wt or T693A, and stimulated or not with FGF10 for 40 or 60 min. showed very good reproducibility for cells expressing siRNA resistant EGFR (Pearson correlation coefficient higher than or equal to 0.85) and very poor correlation for cells expressing EGFR vs. EGFR mutant (Pearson correlation coefficient smaller than 0.2). (H) Cell number increase in T47D cells depleted of EGFR by siRNA, transfected with wt or T693A, and stimulated or not with FGF10 for 72 h. Data represent n=4 experiments. Significance was calculated by one-way ANOVA with Tukey test p=<0.05 *, <0.01 **, <0.001 ***. (I) Lysates from T47D or BT20 depleted of EGFR or not, transfected with wt or T693A, and collected after 24 and 72 hours were immunoblotted with the indicated antibodies.
